# Serotonin modulates learning content-specific neuroplasticity of functional brain networks

**DOI:** 10.1101/2021.01.15.426779

**Authors:** Manfred Klöbl, René Seiger, Thomas Vanicek, Patricia Handschuh, Murray Bruce Reed, Benjamin Spurny, Vera Ritter, Godber Mathis Godbersen, Gregor Gryglewski, Christoph Kraus, Andreas Hahn, Rupert Lanzenberger

## Abstract

Learning-induced neuroplastic changes, further modulated by content and setting, are mirrored in brain functional connectivity (FC). In animal models, serotonergic agents were shown to facilitate neuroplasticity. This is especially prominent during emotional relearning, such as fear extinction, which may translate to clinical improvements in human patients. To investigate this assumption, 99 healthy subjects underwent six weeks of emotional or non-emotional learning and subsequent relearning. Resting-state functional magnetic resonance imaging was performed before and after the learning phases to investigate changes in FC. During relearning, subjects received either a daily dose of the selective serotonin reuptake inhibitor (SSRI) escitalopram or placebo. Escitalopram intake modulated FC changes in a network comprising Broca’s area, the medial prefrontal cortex, the right inferior temporal and left lingual gyrus. More specifically, escitalopram increased the bidirectional connectivity between medial prefrontal cortex and lingual gyrus for non-emotional and additionally the connectivity from medial prefrontal cortex to Broca’s area for emotional relearning. The context-dependence of these effects supports the assumption that SSRIs in clinical practice might improve neuroplasticity rather than psychiatric symptoms per se. Correlations with learning behavior further point towards a relationship with extinction processes in relearning. These results demonstrate that escitalopram intake during relearning results in content-dependent network adaptations and support the conclusion that enhanced neuroplasticity might be the major underlying mechanism also in humans. Beyond expanding the complexities of learning, these findings emphasize the influence of external factors on serotonin-facilitated neuroplasticity of the human brain.

## Introduction

Learning constitutes an evolutionary indispensable process allowing for adjustment to an ever-changing environment. It is accompanied by adaptations in structure and function of the brain, reflected in changes of gray and white matter morphology (Taubert et al., 2012; Valkanova et al., 2014), structural (Sampaio-Baptista and Johansen-Berg, 2017) and functional connectivity (FC) (Guerra-Carrillo et al., 2014). The ability to learn, and thus, neuroplasticity per se, is modulated by the individual’s mental condition (Ehlers, 2012; Taylor Tavares et al., 2008) and the information acquired (Delon-Martin et al., 2013; Draganski et al., 2004; Draganski et al., 2006; Hyde et al., 2009; Maguire et al., 2000). The former is prominently affected in neurological (Filoteo et al., 2007; Grober et al., 2019; Schraegle et al., 2016; Vicari et al., 2005) and psychiatric disorders (Hartmann-Riemer et al., 2017; Marin et al., 2017; Taylor Tavares et al., 2008). A crucial mediating neurotransmitter in these processes is serotonin, which plays a major role in structural (re)modeling of the brain (Daubert and Condron, 2010; Gaspar et al., 2003) and consequently in the pathophysiology and treatment of many psychiatric conditions (e.g., major depression (Kraus et al., 2017), obsessive-compulsive disorder (Fineberg et al., 2012), generalized anxiety disorder (Goodman et al., 2005)). Furthermore, pharmacological modulation of the serotonin system using selective serotonin reuptake inhibitors (SSRIs) was shown to counteract learning deficits in temporal lobe epilepsy (Barkas et al., 2012) but also interact with other neurotransmitter systems (Spurny et al., 2021).

Beyond its involvement in learning processes, serotonin plays a role in extinction and relearning. This is well established in animals (Furr et al., 2012; Lapiz-Bluhm et al., 2009; Masaki et al., 2006) but much less so in humans. A noteworthy finding is faster fear extinction after a two-week treatment with the SSRI escitalopram compared to placebo in healthy subjects (Bui et al., 2013). This effect is suspected to be linked to a positivity bias induced by SSRIs (Harmer and Cowen, 2013; Pringle et al., 2011). Interestingly, such changes were also reported already after a single SSRI dose. Acute application was shown to enhance the recognition of emotional faces (Browning et al., 2007; Harmer et al., 2003) and induce FC changes predictive of treatment response within hours (Klöbl et al., 2020a). Also the acute effects of SSRIs on connectivity depend on the individual mental condition (Dutta et al., 2019). Considering the minimum of one week that is needed for an antidepressant effect (Taylor et al., 2006), these acute findings suggest that serotonin-modulated neuroplasticity facilitates but not necessarily implies improvements in depressive symptoms (Alboni et al., 2017). Thus, serotonergic pharmacological agents can induce widespread and substantial alterations of the brain FC (Arnone et al., 2018; Klaassens et al., 2015; Schaefer et al., 2014; Schrantee et al., 2018).

Since learning induces task-dependent functional network adaptations (Horga et al., 2015; Kang et al., 2018; Lefebvre et al., 2017; Woolley et al., 2015; Zhao et al., 2019), FC provides a convenient surrogate for neuroplasticity. However, several aspects remain unknown. These concern the interactions between learning content and setting (learning vs. relearning) as well as the role of serotonin, especially how the effect of serotonergic agents on emotional relearning (Bui et al., 2013) relates to neuroplastic changes.

In order to map these learning-dependent network adaptations under SSRI intake, neuroplastic changes in FC after learning and relearning were investigated. A dedicated learning paradigm and online training platform were specifically developed for this purpose, comprising an emotional (matching face pairs) and a non-emotional (matching Chinese characters to unrelated German nouns) condition. To assess the modulatory effects of serotonin on neuroplastic changes, participants received escitalopram or placebo during the relearning phase. Resulting from this design, functional network adaptations depending on the learning content and setting were expected with stronger effects of escitalopram on emotional compared to non-emotional relearning. These adaptations were further assumed to correlate with the individual learning behavior.

## Materials and Methods

The study was conducted according to the Declaration of Helsinki including all current revisions and the good scientific practice guidelines of the Medical University of Vienna. The protocol was approved by the institutional review board (EK Nr.: 1739/2016) and the study was registered at clinicaltrials.gov (NCT02753738).

### Study design

The overall study followed a randomized, double-blind, placebo-controlled longitudinal design. Three MRI examinations with 21 days of (re-)learning between each session were conducted (i.e., the MRIs were performed on the 1^st^, 22^nd^ and 43^rd^ day). For a subsample, a test-retest scan was performed 21 days before the baseline, to mitigate the chance of misinterpreting time-as learning-related changes. The subjects were randomized upon recruitment to one of four groups learning to match either Chinese characters to random German nouns or faces to faces and subsequently relearn new associations while receiving placebo or escitalopram (Figure 1A). During the latter phase, the previous associations (character-noun / face-face) were shuffled and had to be relearned following the same time schedule. The study medication consisting of a daily oral dose of 10 mg escitalopram (Cipralex; Lundbeck A/S, Copenhagen, Denmark; provided by the pharmacy of the Medical University of Vienna) or placebo. To monitor the proper intake, the escitalopram blood plasma levels were assessed around day 7, 14 and 21 of the relearning phase.

**Figure 1:**
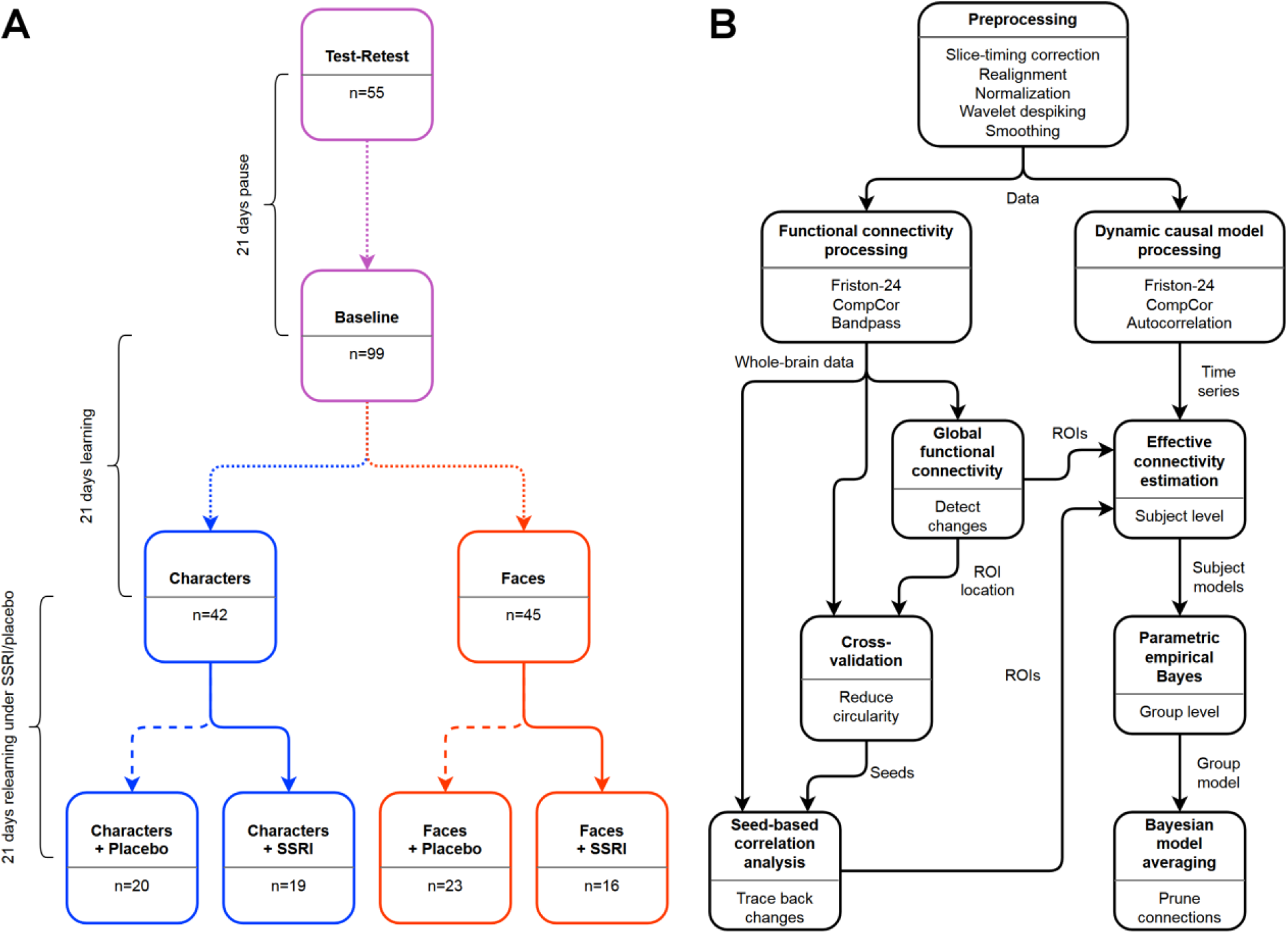
Study workflows. A: Study overview with the number of measurements included at each time point for each group. B: Analysis workflow from the raw data to the final effective connectivity analysis. SSRI: Selective serotonin reuptake inhibitor. ROI: region of interest.

### Learning paradigm

Throughout the course of the study, the subjects had to perform an association-learning task with emotional (face pairs) or non-emotional (Chinese characters – German nouns) content. In both cases they had to learn 200 pairs of images via a daily online training at home (i.e., 21 learning and 21 relearning sessions). Each session contained a pseudorandom selection of 52 image pairs (i.e., the same sequence for all participants). These were presented sequentially for 5 s each. After the training, a pseudorandom selection of 52 images out of all previously seen had to be matched to the correct counterpart without time limit. No feedback was given to keep learning and retrieval strictly separated. All pseudorandomizations were conducted with replacement. The subjects were given personal credentials for the online learning platform and instructed to complete one session per day at approximately the same time. In case sessions were missed, they could be done on the next day. However, subjects were excluded in case of generally irregular learning. During each MRI session, learning and retrieval tasks similar to those on the online platform were performed in the scanner (Reed et al., 2021). In order to minimize additional influences on neuroplasticity, subjects were told not to travel during their participation in the study.

### Participants

In total, 138 healthy volunteers were recruited using advertisements at message boards on the campus of the Medical University and General Hospital of Vienna as well as in libraries, pharmacies and local supermarkets. Inclusion criteria comprised general health based on medical history, physical and psychiatric examination (structured clinical interview (SCID I) for DSM-IV), being 18 to 55 years of age, right-handedness, not smoking and signing the informed consent form. Subjects were excluded in case of psychiatric or neurologic conditions (also in first-degree relatives), MRI contraindications and knowledge of Mandarin, Cantonese or Japanese, positive drug-urine tests, not complying with the study schedule, reported side effects possibly related to the study medication, technical issues and structural anomalies or upon their own request. The distributions of sex, participation in the test-retest session and highest finished level of education between groups were tested using Fisher’s exact (SPSS 25, IBM, Amonk, New York) and that of age with a Kruskal-Wallis test (MATLAB 2018b, Natick, Massachusetts, as all other statistics; all two-sided).

### Modelling the learning behavior

Since the subjects saw only parts of the overall learning content in each session, the respective training results followed a u-shaped curve (Spurny et al., 2020). To correct for this effect, the raw retrieval success was scaled by the relative number of pairs that had already been seen. Weighting the sessions in the modelling process compensated for an overestimation of the training results if two sessions were conducted temporally closer together and an underestimation if further apart (equations (2) and (1)):

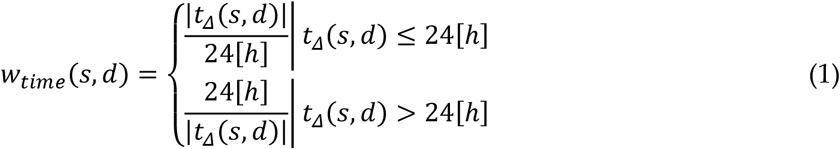

with

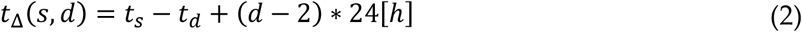

Here, *s* denotes the current and *d* a previous session, *t*_Δ_ is the time difference between expected and actual learning time. Linear discounting was used to reduce the influence of earlier learning times. The total weight *w* for each session was calculated as dot product of the time and discount weights (3).

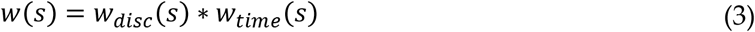

with the discounting weight

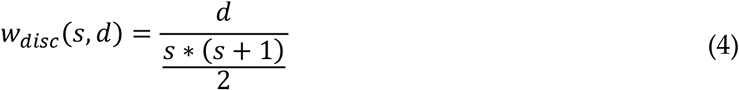

To keep the weighting *w*_*time*_ for a specific session causal, only previous learning times were taken into account, i.e., *d* < *s* in (1) to (4). Using MATLAB, an exponential (5) and a hyperbolic model (6) were fit for each learning phase per subject

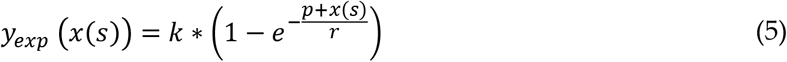

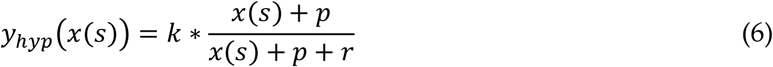

where *x* is the adjusted training success, *k* the “learning capacity” (theoretical maximum of pairs that can be memorized), *r* the “learning rate” (determining how fast k is approached, lower values meaning steeper increase) and *p* the “previous knowledge” from the varying in-scanner session (Anzanello and Fogliatto, 2011). Due to equal complexity, the models were compared by a paired t-test over the Fisher-transformed model fits (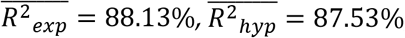, t_164_ = 3.97, p = 1.1E-4 two-sided; the exponential model was preferred). The integral of the fitted learning curve (7) from day 1 to 21 was used to calculate the overall “performance” *Y* (adjusted fraction of correctly retrieved pairs) corrected for irregularities in learning. For further statistical analyses, *Y* was rescaled and Fisher-z-transformed to an unbound distribution (8) (Klöbl et al., 2020b).

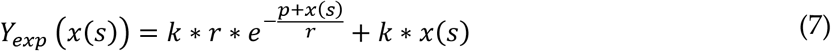

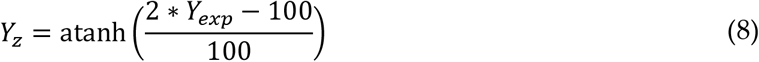

For behavioral analysis, the learning capacity and rate, as well as the adjusted performance were tested for interaction and main effects of the fixed factors “group”, “substance” and “phase” (learning, relearning; as opposed to the three measurements) using linear mixed effects models (LMEs) with a random intercept per subject and an additional random “phase” slope for performance. Covariance structures and random factors were chosen as to minimize the Akaike information criterion. Analyses of variance (ANOVAs) were used to further investigate the effect of “substance” on relearning alone. The learning capacity was rank- and the rate log-transformed as indicated by the residual plots (Conover and Iman, 1981). Values were excluded as outliers if located further than three standard deviations from the mean after transformation. Correlations between the parameters were investigated via partial Spearman correlation, corrected for repetitions over subjects, “group”, “phase” and “substance” conditions. Multiplicity was controlled for using the Sidak correction. All inferences were two-sided.

### MRI acquisition and processing

The RS data was recorded using a Siemens Prisma 3T scanner (Siemens, Erlangen, Germany) equipped with a 64-channel head coil before the in-scanner learning with the following parameters: gradient echo, echo-planar imaging, TE/TR = 30/2050 ms, GRAPPA 2, 210 x 210 mm field of view, 100 x 100 pixel in-plane resolution, 35 axial slices of 2.8 mm (25% gap), flip angle 90°, oriented parallel to the anterior-posterior commissure line, 201 frames (6.9 min). The subjects were instructed to keep their eyes open, lie still and let their mind wander.

The data was preprocessed primarily using Statistical Parametric Mapping, version 12 (SPM12) and custom MATLAB 2018b scripts. Slice-timing correction was performed to the temporally middle slice, followed by two-pass realignment. Images were normalized to the standard space defined by the Montreal Neurological Institute (MNI) and a custom brain mask was applied. The BrainWavelet toolbox (Patel et al., 2014) was used for nonlinear artifact correction with the parameters “chain search” set to “harsh” and “threshold” to “20” to adjust for the application to unsmoothed data and GRAPPA acceleration. The images were then gray-matter-masked and smoothed with a Gaussian kernel of 8 mm full-width at half-maximum.

### Whole-brain functional connectivity analysis

Network changes were identified in a step-wise approach: First, changes in global functional connectivity (GFC) were calculated as a measure of brain-wide connectedness since SSRIs were shown to induce widespread changes thereof (Klaassens et al., 2015; Schaefer et al., 2014). Second, the specific connections underlying these changes were tracked using seed-based correlation analyses allowing for identifying the origins of the GFC changes (Tagliazucchi et al., 2016). Third, spectral dynamic causal models (DCMs) were constructed to infer the directionality of the connections in the network discovered in step 2 (Friston et al., 2016).

Nuisance regression was performed utilizing the Friston-24 model (Friston et al., 1996), an adapted version of the CompCor method with an automated scree approach (Klöbl et al., 2020b) and sine/cosine terms limiting the passband to 0.01-0.10 Hz (Hallquist et al., 2013). GFC maps were calculated by correlation with the standardized average gray matter signal after applying a group mask, which is a parsimonious equivalent of the average correlation to all voxels (Saad et al., 2013). See Figure 1B for a flowchart of the analysis.

The GFC maps were Fisher-transformed and entered into a flexible factorial 2nd-level model in SPM. The model included factors for “group”, “substance” and “measurement” and the results were familywise-error-corrected to p_Cluster_ ≤ 0.025 (two inverse one-sided contrasts, as implemented in SPM12 (Chen et al., 2019)) at peak- or cluster-level (primary peak-level threshold p = 0.001). The interaction effects were estimated and post-hoc comparisons adjusted again using the Sidak method.

For deducing which regions had the strongest influence on the changes in GFC, the analysis was repeated in a 10-fold cross-validation (the clusters were visually identified disregarding significance due to the reduced sample size). This way, the inherent circularity of inferences on the results is reduced. The first temporal eigenvariate from each significant cluster was extracted via the MarsBaR toolbox and used for a seed-based correlation analysis (SBCA). The Fisher-transformed SBCA maps were fed into the same model as used for the GFC data. Results were corrected for the number of seeds and post-hoc comparisons using the Sidak method.

### DCM analysis

For the DCMs, the smoothed data was reprocessed using the 1st-level GLM in SPM12 again correcting for the Friston-24 and CompCor regressors (Esménio et al., 2019). Autocorrelation was set to “FAST” (Olszowy et al., 2019). The first temporal eigenvariates of clusters from the GFC and the SBCA analyses surviving multiplicity correction were extracted from the data preprocessed for the DCM analysis. With these, fully connected linear spectral two-state DCMs were estimated. The parametric empirical Bayes (PEB) framework in SPM12 was used for group inference. A flat model was compared to a hierarchical PEB-of-PEBs approach in terms of free energy using only the subjects that completed all three scans from baseline to relearning. The former model has the advantage to allow for inclusion of partially available datasets whereas the latter can better account for within-subject effects by first creating PEB models for the individual subjects which are then fed into a group analysis. Since the flat model was favored the terms of free energy, this model was employed. To control for potential purely temporal effects, the test-retest scans were included as additional measurement and a correction factor for subjects that participated in these. Bayesian model averaging (Friston et al., 2016) was finally utilized to prune connections with high uncertainty.

In order to assess the dependencies between learning behavior and changes in effective connectivity, a PEB-of-PEBs model was set up with the differences between the scans after to before the respective phases on the lower level. The PEB-of-PEBs approach was here used to account for parameter certainty when calculating the difference. Since varying results for the different conditions were expected, interactions of the single learning parameters with “phase”, “group” and “substance” were investigated. The learning parameters were transformed as before. No outliers were excluded at this stage. Final Bayesian model reduction was applied as above. Unless otherwise mentioned, standard settings were used in the DCM analysis.

## Results

Out of 138 subjects recruited, 99 subjects participated in the first MRI session. Of those, 87 completed the second and 78 also the third MRI scan. Additionally, 55 participants partook in the initial test-retest session (Figure 1A). The subjects that at least completed the baseline MRI were 25 ± 5 years old (median ± interquartile range) and comprised 56 women and 43 men. There were no significant group differences regarding age, sex, highest finished level of education or participation proportions in the test-retest session (all *p* > 0.2).

### Learning behavior

Figure 2A shows example learning curves and fitting details for two subjects. An overview of the parameter distributions is given in Figure 2B. The significant results are presented in Table 1. The models show that learning was more difficult for the “faces” condition and during the “relearning” phase. The relationships between the parameters imply that higher learning capacities were reached later and drove performance. No significant behavioral interactions or substance effects were found, indicating no influence of escitalopram on modeled learning behavior.

**Figure 2:**
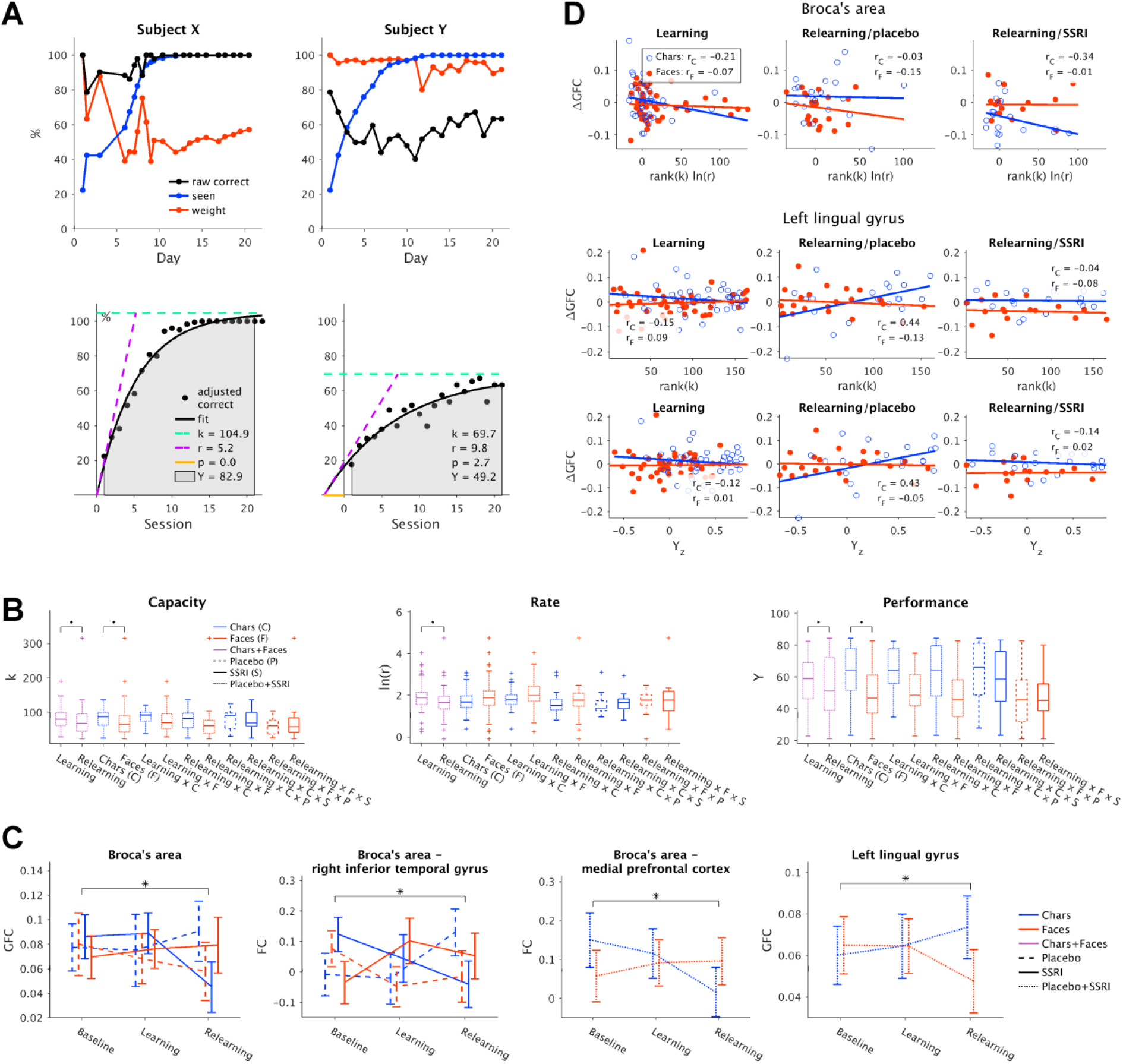
Overview of learning and functional connectivity results. A: Example learning data and model fits over 21 days for two subjects. The percentages of correct answers, cumulative image pairs and model fitting weights calculated from the regularity of learning are shown in the upper row. The lower row shows the adjusted learning curves, model fits and derived learning parameters. B: Box plots of the learning parameters for the single conditions (the exponential learning rate was log-transformed for better visibility). Center line: median, box limits: quartiles, whiskers: most extreme non-outlier points, points: outliers further than 1.5 times the interquartile range from the quartiles. C: Means and 95%-confidence intervals of all available scans for the significant global (GFC) and inferred functional connectivity (FC) differences between sessions and conditions. D: Influences of the learning parameters (transformed due to outliers and skewed distributions) on the changes in GFC of 78 subjects. The different correlations and overlaid slopes demonstrate a varying influence of the learning parameters on GFC ^*^ indicates significant tested differences.

**Table 1:** Significant influences of experimental variables on learning parameters “capacity” k, “rate” r and “performance” Y and correlations between them (N = 87). Linear mixed effects models were used for comparisons between conditions on the transformed parameters. Partial Spearman correlations were corrected for repetitions over subjects and the influence of learning phase and content and substance condition. Confidence intervals (CI) and p-values are adjusted for the three comparisons.^1^ k was rank-transformed: Estimate and CI were derived from the average increase in k per rank. ^2^ r was log-transformed: Estimate and CI represent fractions of geometric means. ^3^ Y was rescaled and Fisher-transformed: Estimate and CI are given for untransformed data to ease interpretability. DoF: degrees of freedom.

| Parameter | Contrast | Estimate | 98.3%-CI | DoF | t-value | p-value |
| --- | --- | --- | --- | --- | --- | --- |
| k | faces-chars | -16.15<sup>1</sup> | -29.74, -2.57 | 161 | -2.87 | 0.0140 |
| k | relearn.-learn. | -11.17<sup>1</sup> | -16.32, -6.02 | 161 | -5.23 | 1.5E-6 |
| r | relearn.-learn. | 0.77<sup>2</sup> | 0.65, 0.92 | 158 | -3.53 | 0.0016 |
| Y | faces-chars | -14.90<sup>3</sup> | -21.97, -7.83 | 161 | -5.15 | 2.2E-6 |
| Y | relearn.-learn. | -2.98<sup>3</sup> | -5.40, -0.57 | 161 | -2.58 | 0.0318 |
| Parameter 1 | Parameter 2 | Correlation | 98.3%-CI | DoF | p-value |  |
| k | r | 0.54 | 0.36, 0.69 | 84 | 2.29E-7 |  |
| k | Y | 0.53 | 0.34, 0.68 | 84 | 8.26E-7 |  |

### Whole-brain functional connectivity

A significant interaction effect in GFC between “group”, “substance” and “measurement” (post-relearning compared to baseline) was found in Broca’s area (BA; Table 2, Figure 2C), showing a marked decrease during relearning of character-noun associations under escitalopram. A second interaction of “group” and “measurement” but without influence of “substance” was found after relearning in the left lingual gyrus (lLG) with an increase for characters and a decrease for faces.

**Table 2:**
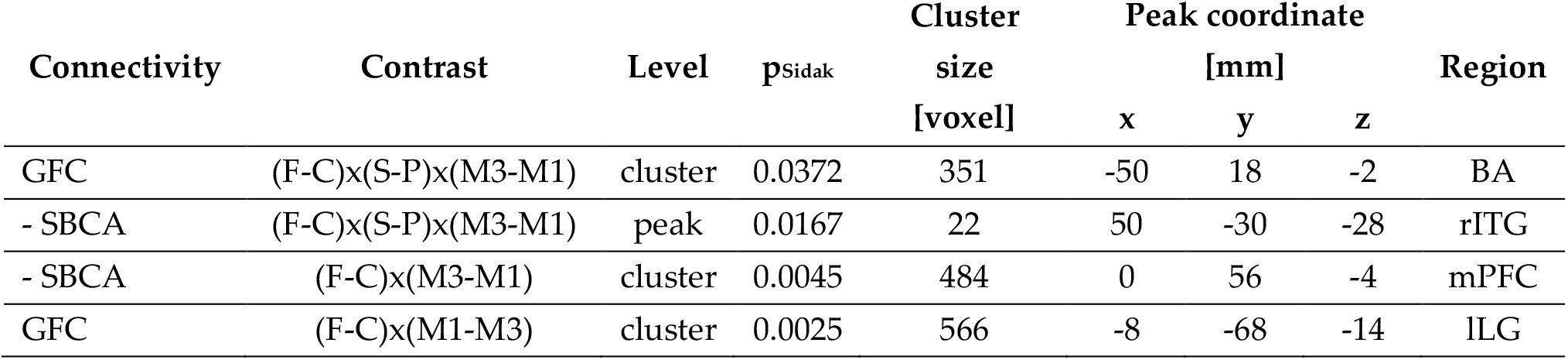
Whole-brain family-wise-error-corrected cluster-level results for the global functional connectivity (GFC) and the subsequent seed-based correlation analyses (SBCA) listed below. Factors: “group” (faces “F” / Chinese characters “C”), “substance” (placebo “P” / SSRI “S”) and “measurement” (“M1…3”). BA: Broca’s area, rITG, right inferior temporal gyrus, mPFC: medial prefrontal cortex, lLG: left lingual gyrus.

Re-estimating the statistical model above with the SBCA maps revealed changes in the right inferior temporal gyrus (rITG) and the medial prefrontal cortex (mPFC) for the BA. No SBCA results for the lLG seed survived multiplicity correction. As the placebo and SSRI groups were not treated differently between the baseline and the post-learning session, any such temporal variations arouse due to external factors not examined in this study.

Since both GFC results suggest an influence of learning in general rather than relearning alone, the effect of behavior on the GFC changes was further investigated on an exploratory basis. For this, the median values for the significant regions were extracted using the MarsBaR toolbox and changes in GFC modeled via LMEs (p < 0.05, two-sided) depending on the interaction of “group”, “substance”, “phase” and the transformed learning parameters. Associations with learning parameters substantiated the assumption for BA by a significant condition-dependent influence of capacity and rate (p = 0.0249, t_130_ = 2.27, N = 87). Further, the GFC change in the lLG could be similarly modeled by performance (p = 0.0133, t_149_ = 2.50, N = 87) or capacity (p = 0.0142, t_149_ = 2.48, N = 87; see Figure 2D). These relationships indicate that the influence of learning on GFC changes with content, setting and substance. No significant changes in GFC between directly consecutive scans (including test-retest) were found.

### Effective connectivity

The DCMs explained 86.51 ± 3.96% of the single-subject variance (median ± interquartile range) indicating an adequate fit. Figure 3 shows the learning-specific effects of the final Bayesian model averaging after PEB inference. Connections generally increased from the test-retest to the baseline session and decreased throughout learning.

**Figure 3:**
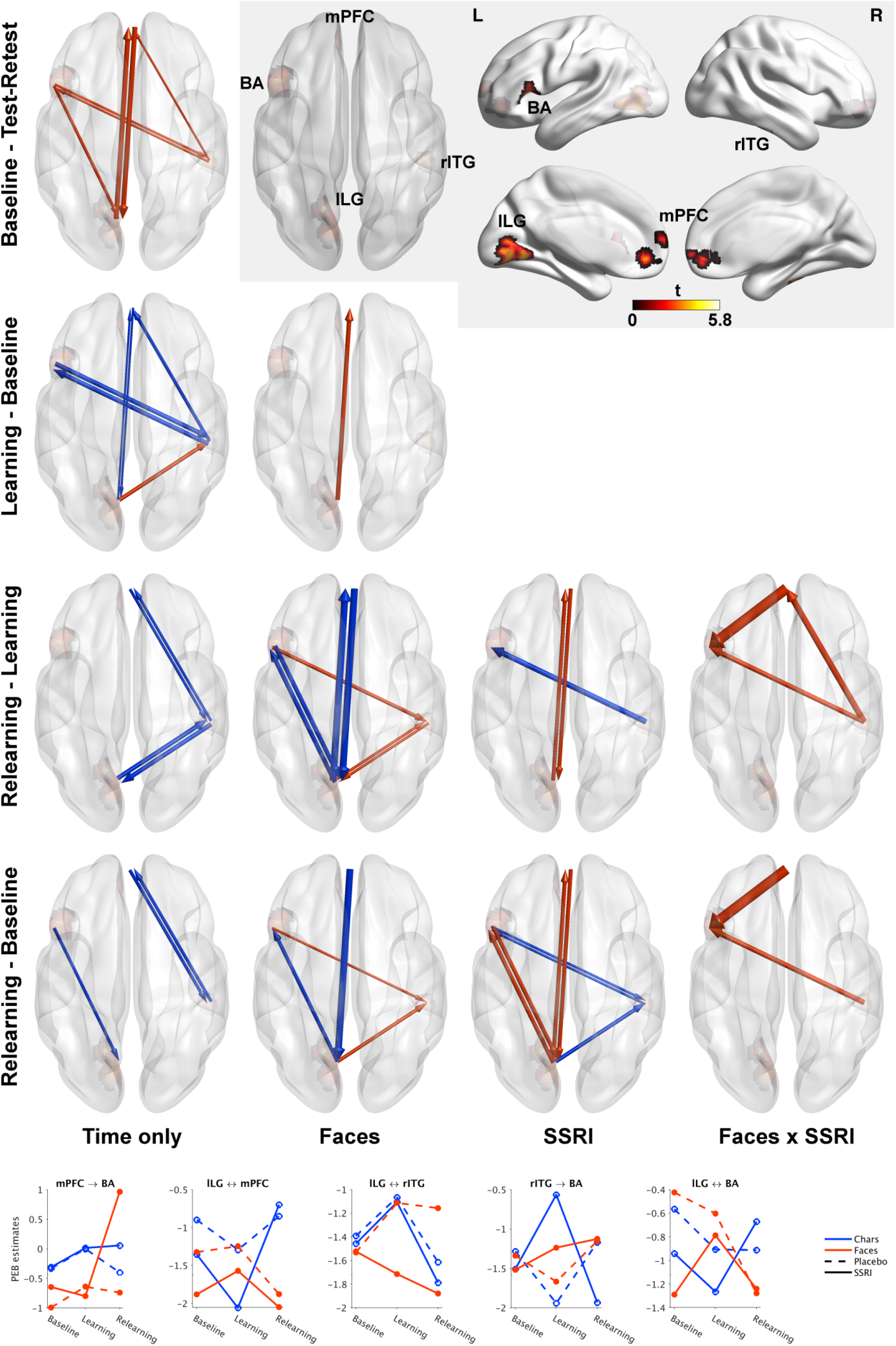
Effective connectivity time and time-related interaction effects relative to the reference conditions (learning Chinese characters, placebo application). Increases are indicated in red, decreases in blue, line thickness represents the expected value of change in effective connectivity. The shaded inlay shows the cluster positions and t-values of the preceding global and seed-based functional connectivity analyses that were used to define the regions of interest (primary threshold ***p*** **≤ 0.001** – equivalent ***t*** **≥ 3.15**, ***p***_***cluster***_**≤0.05**) Effects with a posterior probability > 99% are shown. The temporal changes of selected estimates of the preceding parametric empirical Bayes (PEB) analysis are displayed below (estimates were averaged where multiple connections are shown at once). Here, the right three graphs also show unsystematic effects within the subgroups from baseline to post-learning. BA: Broca’s area, tITG: right inferior temporal gyrus, mPFC: medial prefrontal cortex, lLG: left lingual gyrus, SSRI selective serotonin reuptake inhibitor. Figure created with BrainNet Viewer 1.7 (Xia et al., 2013).

Emotional relearning under escitalopram led to a drastic increase in the otherwise comparably stable connectivity from the mPFC to BA. During learning, the bidirectional connections between mPFC and lLG showed an increase for emotional and a decrease for non-emotional content. This was followed by a marked decrease during emotional relearning under placebo as well as an increase during non-emotional relearning under escitalopram. A strong decrease in connectivity during non-emotional relearning was observed between the lLG and the rITG. However, the different time courses of the “faces” subgroups from baseline to post-learning do not allow for the interpretation of prolonged learning effects. Similar unsystematic deviations over time affect the connections between rITG / lLG and BA.

### Relationships between learning behavior and connectivity changes

In order to allow for conclusions on the influence of learning capacity, rate and performance, the dependence of effective connectivity changes on these parameters was estimated (Figure 4). Correlations for learning per se were mostly weak compared to the changes related to the experimental conditions.

**Figure 4:**
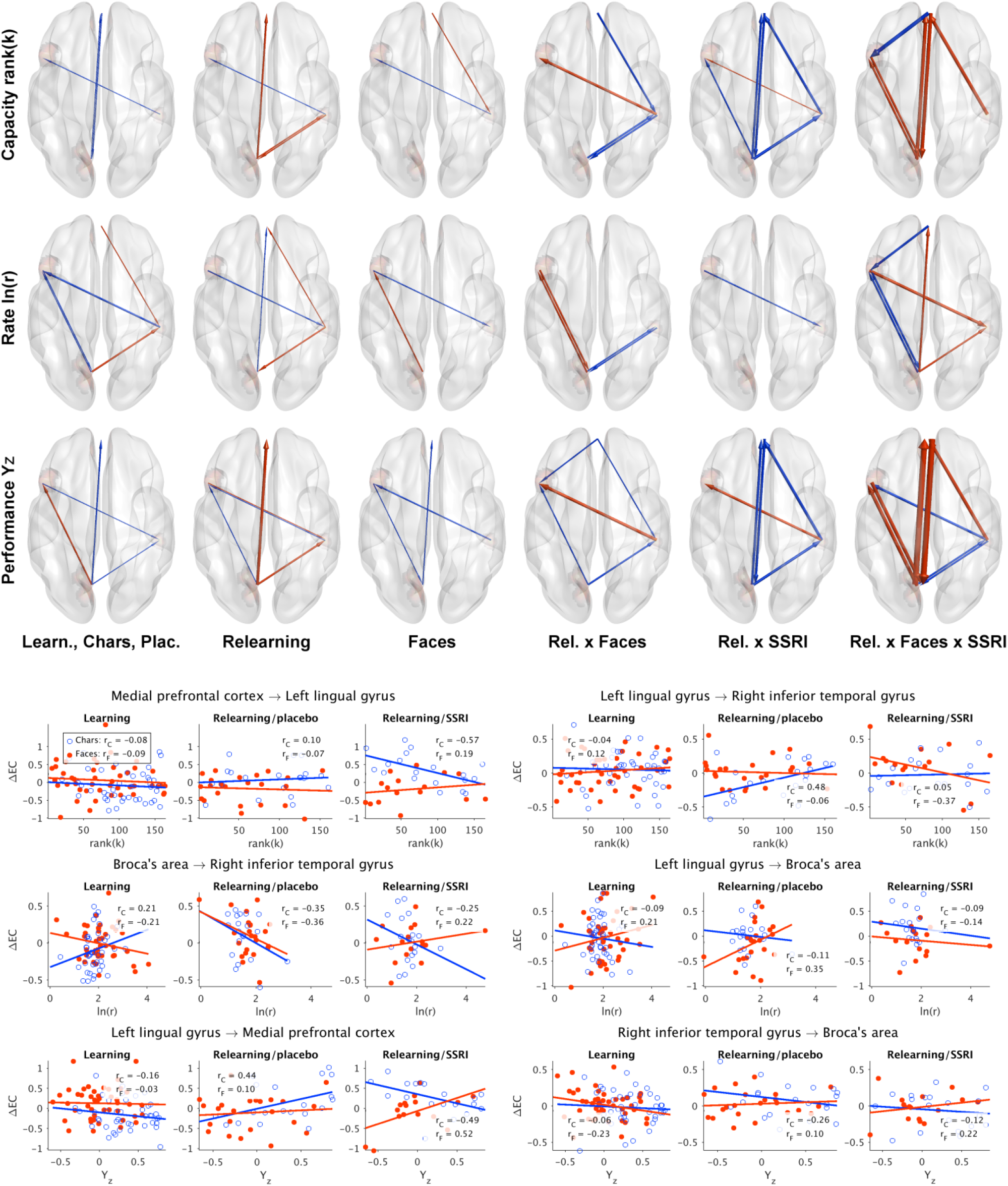
Correlations between changes in effective connectivity (ΔEC) and the learning parameters capacity, rate and performance. Strengthened correlations are indicated in red, weakened in blue, line thickness represents the expected value of the relationship change. The initial learning phase, matching nouns to Chinese characters and placebo were used as reference conditions (left column). Effects with a posterior probability > 99% are shown. The correlations of selected effective connection changes with learning behavior for the 78 subjects are detailed below. Different relationships are visible for learning / relearning, the “faces” and “characters” groups and pharmacological modulation via the selective serotonin reuptake inhibitor (SSRI) escitalopram. Learning capacity, rate and performance were rank-, log- and Fisher-transformed beforehand to avoid skewed distributions and outliers. Figure created with BrainNet Viewer 1.7 (Xia et al., 2013).

Learning capacity and performance showed similar dependencies including negative correlations with the connectivity changes between lLG and mPFC for non-emotional learning, which increased for relearning. The intake of escitalopram during relearning led to a strong positive dependency for emotional, and negative for non-emotional learning content. The connections between lLG and rITG showed similar correlations but were positive for relearning character-noun associations under placebo and negative for face pairs under escitalopram. Variations lower in magnitude were also found for the dependence of the rITG-BA connectivity changes. The connection from BA to the rITG was anti-correlated with learning rate. This relationship was further enhanced during relearning and for the “faces” group. On the contrary, the connections between the lLG and BA showed an increased positive relationship with the rate for emotional (re-)learning, which is suppressed by the application of escitalopram.

## Discussion

A functional brain network sensitive to the interaction of learning content (emotional, non-emotional), setting (learning, relearning) and serotonergic modulation was identified. Contrary to the initial assumption, the application of the SSRI escitalopram also had a limited influence on the connectivity changes induced by non-emotional relearning.

### Context-dependent communication between medial prefrontal cortex and lingual gyrus

Three weeks of escitalopram intake led to an increase in connectivity between lLG and mPFC for non-emotional relearning, implying stronger integrative processing between these regions. In mice, the chronic application of the SSRIs was found to induce dendritic spine growth in the mPFC (Guirado et al., 2014) and promote plasticity of the visual cortex (Chen et al., 2011; Maya Vetencourt et al., 2008). After acute citalopram administration in humans, an increase in mPFC FC with the dorsolateral prefrontal and posterior cingulate cortex was discovered (Arnone et al., 2018). In contrast, reductions in global (Schaefer et al., 2014) and network-specific FC (Klaassens et al., 2015) were also reported. Given the increases after 21 days of relearning under escitalopram, this points towards content- and regionally specific changes of neuroplasticity.

Citalopram has been previously linked to reduced activation in the lingual gyrus when viewing emotional faces (Henry et al., 2013). The current results imply an accompanying increase in connectivity towards the mPFC when associations with these faces should be relearned. Activity of mPFC and rITG have been related to the facial expression observed in others (Zaki et al., 2012). Both regions are also important for durable memory encoding (Wagner et al., 2016; Wagner et al., 2019). Furthermore, the mPFC plays an explicit role in memorizing emotional faces (Keightley et al., 2011). Thus, conflicting emotional memories could be mirrored in the opposed connectivity changes of mPFC and lLG when learning and relearning faces.

The LG has been linked to extinction learning (Klass et al., 2017; Lissek et al., 2015a; Lissek et al., 2015b) and structural alterations to panic (Pang et al., 2019) and posttraumatic stress disorders (Kunimatsu et al., 2020). Both conditions are suspected to be based on dysfunctional fear learning and extinction. The central role in extinction processes is further backed by the correlation found between LG-mPFC and LG-rITG connectivity changes and relearning capacity and performance. An extinction-related network comprising the mPFC, hippocampus and right amygdala was previously identified in fear conditioning (Lang et al., 2009).

Figure 3 reveals that the connections between mPFC and lLG were weakened when relearning faces and strengthened for characters. The latter effect was markedly potentiated under escitalopram. This differential pattern extends previous results that SSRIs facilitate neuroplasticity not just in general but in a context-specific manner. Considering that the improvement of clinical symptoms in psychiatric disorders may actually be driven by the neuroplastic action of SSRIs (Chiarotti et al., 2017), it seems particularly important to provide a well-designed environment for the treatment of these patients.

Under escitalopram, the connections from lLG to mPFC further showed a distinct relationship to learning performance, with a strong positive association for relearning faces and a negative one for characters. Given the increase in connectivity for the “characters” group under escitalopram, higher performance implies a stronger reduction, also supporting the extinction perspective. Thus, under escitalopram, a decreased connectivity between lLG and mPFC is accompanied by an increased dependency of communication on learning performance. The bidirectionality of the decrease might be related to a serotonergic modulation of affective feedback processes (Rudrauf et al., 2008).

### The role of Broca’s area in learning

BA is involved in numerous aspects of speech (Fujii et al., 2016), including inner speech (Morin and Hamper, 2012), and mnemonic strategies (Love et al., 2006). Even though reading words was shown to lead to electrical activity in BA (Magrassi et al., 2015) no relationship to nouns as linguistic objects was found (Faroqi-Shah et al., 2018). Theories of a topologically distinct representation of nouns in temporal regions (Vigliocco et al., 2011) have also failed to gain meta-analytical support (Crepaldi et al., 2013) making it unlikely that changes in these regions stem from the learning content alone. Under acute tryptophan depletion, BA showed a decrease and the mPFC an increase in activation for frontal-compared to side-viewed faces (Williams et al., 2007), which ascribe BA also a serotonergic modulatable role in emotion processing.

After relearning faces under escitalopram, a strong increase in connectivity from the mPFC towards BA was detected. For the same learning content, a bidirectional decrease in connectivity between BA and lLG was found independent of the drug condition. The enhanced connectivity might indicate serotonergic facilitation of emotional relearning and provides support for the importance of BA in emotion processing (Williams et al., 2007) and the context for neuroplastic changes (Alboni et al., 2017; Chiarotti et al., 2017).

The connectivity changes between the lLG and BA showed a positive dependence on the learning rate r for learning as well as relearning of face pairs under placebo. Since a higher rate corresponds to a flatter slope of the learning curve, subjects with stronger increases in connectivity approached their learning capacity slower. As this finding is accompanied by decreases in dependency on performance, it might be more indicative of slow learning than cognitive reserves. The connectivity changes between lLG and BA when initially learning character-noun associations is positively correlated with performance, which might be expected based on the role of BA in language. In this light, the strong increase of the correlation of lLG-BA connectivity changes and capacity / performance for emotional relearning under escitalopram points towards a facilitating effect of serotonin on learning-related neuroplasticity. The complexity of this relationship is also indicated by the significant interaction of learning capacity and rate on GFC (Figure 2D).

### Modulation along the ventral visual stream

The connections between the lLG and the rITG run along the ventral visual stream (VVS). It generally connects the visual and the inferior temporal cortex and is implicated in object recognition and identification (Goodale and Milner, 1992). The plasticity of this pathway and the effects of its modulation via transcranial direct current stimulation on memory encoding were recently shown (Zhao and Woodman, 2021). The inferior temporal cortex itself is also involved in short-term (Ranganath et al., 2004) and long-term memory (Wagner et al., 2016; Wagner et al., 2019), object naming and identification (Acres et al., 2009).

For relearning, a bidirectional decrease in connectivity between lLG and rITG could be observed for emotional as well as non-emotional content, being much more pronounced for the latter one. A systematic effect of escitalopram on connectivity was not observed. It did, however, modulate the dependency of connectivity changes on capacity / performance. The positive correlation for relearning character-noun associations was not present under escitalopram. The dependency became even negative for face pairs.

The importance of the LG for visual memory is well-established (Bogousslavsky et al., 1987) together with its involvement in facial (Puce et al., 1995) and word form processing (Mechelli et al., 2000; Xiao et al., 2005). Hemispheric differentiation was previously suggested with the lLG being more active during memorizing faces and the right LG during passive viewing (Kozlovskiy et al., 2014). This is also reflected in the current results as increased connectivity from lLG to mPFC after learning to match face pairs. Besides visual memory and processing, the LG and inferior frontal gyrus, where BA is located, were shown to be important for the analysis of novelty and spatial information (Menon et al., 2000). This might explain the differences in effective but not global connectivity between the test-retest and baseline scan. Where the information flow for visual processing is directed from the visual to the temporal cortex, also feedback mechanisms from emotion-related structures were shown (Rudrauf et al., 2008). Moreover, plasticity in the visual cortex was demonstrated following emotional learning (Meaux et al., 2019). Interestingly, the connection from lLG to rITG is the only one showing an increase after initial non-emotional learning. A possible explanation might lie in object identification or memory formation processes related to the rITG. The more wide-spread reductions could be based on the decreasing novelty or repetition suppression (Prčkovska et al., 2017). Serotonergic modulation of the behavioral dependence of connections along the VVS might be based on long-term effects of escitalopram on the rITG (Kaichi et al., 2016) or facilitation of neuroplasticity of the lLG as part of the visual cortex (Chen et al., 2011; Maya Vetencourt et al., 2008).

### Limitations

Despite the comparably large sample, the dropout rate led to slightly imbalanced subgroups. Models allowing for missing values were utilized where possible to mitigate this problem. Caution is needed when interpreting certain results in light of accidental subgroup differences at baseline and after initial learning. The test-retest session performed to differentiate temporal and general learning effects probably had an effect on the identified learning network due to shared processing of novelty. Despite correcting for such effects, the in-depth discussion thus concentrated on the interactions of the experimental conditions which should not be affected so easily. Even though serotonergic modulation of the correlations between connectivity and behavior was found, no direct influence of escitalopram on learning performance itself was detected. However, also previous findings on effects of SSRIs on learning performance were contradictory (Barkas et al., 2012; Chamberlain et al., 2006).

## Conclusion

A learning content, setting sensitive and serotonergic modulated functional brain network and its behavioral correlates were mapped. Three prominent network-specific scenarios were identified: Between the mPFC and the lLG, the intake of escitalopram during relearning potentiated the bidirectional connectivity increase for charcter-noun associations. Further, for relearning face pairs, the directed connection from mPFC to BA was drastically strengthened only under escitalopram. These findings point towards a context-dependent facilitation of neuroplasticity. Lastly, along the VVS, even though the intake of escitalopram did not directly influence the connectivity between lLG and rITG, it did modulate the dependency on learning behavior. These content-dependent changes match the theory that in depression SSRIs improve neuroplasticity rather than mood (Alboni et al., 2017; Chiarotti et al., 2017). This would make patients more susceptible to environmental influences, which ideally provide a setting that supports the therapeutic endeavor.

Moreover, prominent context-dependent correlations of the relearning-induced connectivity changes with performance were found from the lLG to the mPFC, which might be related to extinction of previously learned content. A challenge for future studies addressing the highly complex interactions between learning, network adaptations, serotonergic modulation and behavior will be to adequately control for phenomena with common characteristics, such as the recognition of novelty. Finally, the results on context-dependent neuroplasticity require consideration in treatment studies using serotonergic medication, as they necessitate increased attention towards external factors.

## Data availability

The preprocessed data underlying the analyses in this work is available from the corresponding author upon reasonable request. Due to data protection, we are not able to share the raw data.

## Code availability

In-house code used for the analyses in this work is available from the corresponding author upon reasonable request.

## Acknowledgments

This research was funded in part, by the Austrian Science Fund (FWF) [KLI 516, PI: Rupert Lanzenberger]. For the purpose of open access, the author has applied a CC BY public copyright license to any Author Accepted Manuscript version arising from this submission. This work was further supported by the Medical Imaging Cluster of the Medical University of Vienna, and by the grant „Interdisciplinary translational brain research cluster (ITHC) with highfield MR” from the Federal Ministry of Science, Research and Economy (BMWFW), Austria. Vienna. Manfred Klöbl and Murray Bruce Reed are recipients of a DOC fellowship of the Austrian Academy of Sciences at the Department of Psychiatry and Psychotherapy of the Medical University of Vienna. The funding agencies had no role in conceptualization, design, data collection, analysis, decision to publish, or preparation of the manuscript. We want to thank Dietmar Winkler for providing clinical supervision, Leo Silberbauer, Jakob Unterholzner, Paul Michenthaler, Alim Basaran and Alexander Kautzky for medical support. We also want to express our gratitude towards the diploma students of the Neuroimaging Labs, especially Hannah van Alebeek who was responsible for recruiting a large proportion of the volunteers for the study.

## Author Contributions

Rupert Lanzenberger was the principal investigator and supervised the project supported by René Seiger and Thomas Vanicek. Manfred Klöbl planned and conducted the analysis and wrote the first draft of the manuscript. Murray Bruce Reed, Benjamin Spurny, René Seiger and Manfred Klöbl acquired the magnetic resonance imaging data. The online learning platform was implemented by René Seiger. Andreas Hahn supervised the analysis and provided methodological support. Vera Ritter supported the subject recruiting and management. Gregor Gryglewski, Rupert Lanzenberger and Christoph Kraus wrote the study protocol. Medical support was provided by Thomas Vanicek, Patricia Handschuh and Godber Mathis Godbersen. All authors read and critically revised the manuscript and agreed to the final version.

